# Uncoupling DNA- and RNA-directed DNA methylation at *Rasgrf1*

**DOI:** 10.1101/297465

**Authors:** Erin T. Chu, David H. Taylor, Margaret Hofstedt, Paul D. Soloway

**Author notes:** Current Address: Department of Molecular, Cellular, and Developmental Biology, Yale University, New Haven, CT 06520.

## Abstract

Long noncoding RNAs (lncRNAs) have garnered much attention as possible links between DNA sequence and the protein factors that mediate DNA methylation. However, the mechanisms by which DNA methylation is directed to specific genomic locations remain poorly understood. We previously identified a lncRNA in mouse, the pitRNA, that was implicated in the control of DNA methylation at the imprinted *Rasgrf1* locus. The pitRNA is transcribed in the developing male germline antisense to the differentially methylated region (DMR) that harbors paternal allele methylation, and is driven by a series of tandem repeats that are necessary for imprinted methylation.

MitoPLD, a factor necessary for piRNA biogenesis, both processes piRNAs from the pitRNA, and is necessary for complete methylation at the locus, along with piRNA binding proteins. Using two independent mouse systems where pitRNA transcription is driven by the doxycycline-inducible Tet Operator, we demonstrate that pitRNA transcription across the DMR is insufficient for imprinted methylation, and that the *Rasgrf1* repeats have additional, critical *cis*-acting roles for imparting DNA methylation to *Rasgrf1*, independently of their control of pitRNA transcription. Furthermore, pitRNA overexpression and oocyte loading of pitRNA is insufficient to induce transallelic and transgenerational effects previously reported for *Rasgrf1*. Notably, manipulation of the pitRNA with the *TetOFF* system led to transcriptional perturbations over a broad chromosomal region surrounding the inserted Tet Operator, revealing that the effects of this regulatory tool are not localized to a single target gene.

**AUTHOR SUMMARY:** DNA methylation is a heritable genetic modification known to impact vital biological processes. While the proteins that establish, maintain, and remove DNA methylation are well characterized, the mechanisms by which these proteins are directed to specific genetic sequences are poorly understood. We have previously demonstrated that DNA methylation at the imprinted *Rasgrf1* locus requires a DNA element with a series of tandem repeats. These repeats act as a promoter for a long noncoding RNA, the pitRNA, which is targeted by a small noncoding RNA pathway known to silence viral elements in the male germline via DNA methylation. We queried the sufficiency of the pitRNA to mediate DNA methylation at *Rasgrf1*. We show that, in the absence of the repeats, the pitRNA expression is insufficient to establish imprinted methylation. This work supports a pitRNA-independent mechanism for methylation at *Rasgrf1*, and a critical *cis*-acting role for the tandem repeats separate from their control of pitRNA transcription.

## INTRODUCTION

DNA methylation is essential for appropriate embryonic development. While the *trans*-acting factors required to establish [1,2,3,4,5], maintain [6,7,8,9,10,11], and remove [12,13,14,15] DNA methylation have been identified, little is known of *cis*-acting elements that direct these *trans*-acting factors to specific genomic locations [16,17,18,19,20].

One such *cis* element exists at the imprinted *Rasgrf1* locus. In mouse and rat, *Rasgrf1* is paternally methylated and expressed in the neonatal brain. Imprinted expression in mouse is controlled by a differentially methylated region (DMR) 30 kb upstream of *Rasgrf1* coding sequence, and requires a 1.6 kb stretch of tandem repeats immediately adjacent to the DMR. Targeted deletion of the *Rasgrf1* repeats (*Rasgrf1^tm1^*^PDS^, *tm1*) leads to loss of methylation at the *tm1* DMR in the male germline, and loss of imprinted *Rasgrf1* expression [21]. The repeats also play a role in the maintenance and spreading of DMR methylation in the embryonic somatic lineage after fertilization, though they are dispensable past the epiblast stage [22,23].

Our lab previously characterized a long noncoding RNA (lncRNA), the pitRNA, which is driven by the *Rasgrf1* repeats and is expressed in the embryonic male gonad [24]. LncRNAs, with their ability to recruit and bind effector proteins [25,26], represent a molecular class that could bridge the gap between the protein effectors of local epigenetic states if they recruit the effectors while being transcribed. Indeed, lncRNAs have been implicated in diverse biological processes, and have been proposed to modulate gene expression via a number of mechanisms including recruitment of histone modification complexes [27,28], transcriptional interference [29], and enhancer regulation [30].

Using the *Rasgrf1* repeats as a promoter, the pitRNA is transcribed antisense to the DMR, spanning an RMER4B element, an LTR-type retrotransposon. The repeats and RMER4B element are conserved at the *Rasgrf1* DMR in species where *Rasgrf1* is imprinted [31]. The pitRNA is processed into secondary piRNAs by the piRNA pathway, which is required for DNA methylation and transcriptional silencing of retrotransposons [32], and also for full methylation at the *Rasgrf1* DMR [24]. Given the apparent importance of the pitRNA and piRNA pathway in controlling methylation at *Rasgrf1*, we hypothesized that aberrant expression of the pitRNA could explain transallelic and transgenerational effects previously reported at *Rasgrf1* [33]. Indeed, aberrations in non-coding expression have been associated with such effects in other systems [34,35,36].

More recently, our lab targeted the *Wnt1* locus, inserting the *Rasgrf1* repeats and DMR between the *Wnt1* coding sequence and its annotated enhancer (*Wnt1*^*DR*^). We found that when paternally transmitted, the *Wnt1*^*DR*^ allele was methylated, recapitulating patterns of imprinted methylation found at *Rasgrf1*. However, pitRNA expression from the *Wnt1*^*DR*^ was extremely low (less than 2% of the pitRNA expressed from the endogenous locus) [37]. These data suggested that the *Rasgrf1* repeats could impart methylation to their associated DMR independent of robust pitRNA expression.

None of the systems described above have uncoupled the pitRNA from the *Rasgrf1* repeats to ascertain necessity or sufficiency of either individual element for methylation in *cis* at the endogenous *Rasgrf1* locus. Here, we directly queried the sufficiency of the pitRNA to establish methylation at *Rasgrf1* using a targeted mutation in mouse, *Rasgrf1*^tm5.1PDS^ (*tm5.1*) where the *Rasgrf1* repeats were replaced by the Tet Operator. This enabled inducible control of pitRNA expression through combination of *tm5.1* with one of two transactivating proteins: *TetON*, which binds the Tet Operator and drives pitRNA expression in the presence of doxycycline [38]; and *TetOFF*, which binds the Tet Operator and drives pitRNA expression in the absence of doxycycline [39,40].

We found that induction of the pitRNA at physiologic levels in male gonocytes was insufficient to impart methylation to the *tm5.1* DMR, revealing a critical role for the repeats in methylation control, independent of their regulation of pitRNA transcription, consistent with our findings with *Wnt1*^*DR*^. Using *tm5.1* as well as a transgenic allele, TetO∆^Tg^, we also determined that the pitRNA overexpression was insufficient to induce transgenerational or transallelic effects on *Rasgrf1* expression or methylation. Finally, in addition to enabling control of pitRNA expression as designed, *TetOFF* transactivation of *tm5.1* activated transcription across a broad chromatin domain previously shown to exhibit interactions, and that activation was not confined to the target sequences at the DMR. Our data identify a role for *Rasgrf1* repeats as a *cis*-element directing DNA methylation, independently of the pitRNA it drives. Furthermore, we show that expression patterns controlled by engineered Tet repressor proteins can be exerted over large regions of the genome.

## RESULTS

### Generation of *Rasgrf1^tm5.0^*^PDS^

We successfully generated the targeted mutant *Rasgrf1*^tm5.1PDS^ (*tm5.1*), where the endogenous repeats were replaced with the Tet Operator (Fig 1a). Allelic structure was validated by Southern blot, and Sanger sequencing of PCR products that spanned the vector ends and the endogenous sequence at the target locus, as well as by copy number qPCR (Fig S1a-d).

**FIGURE 1.**
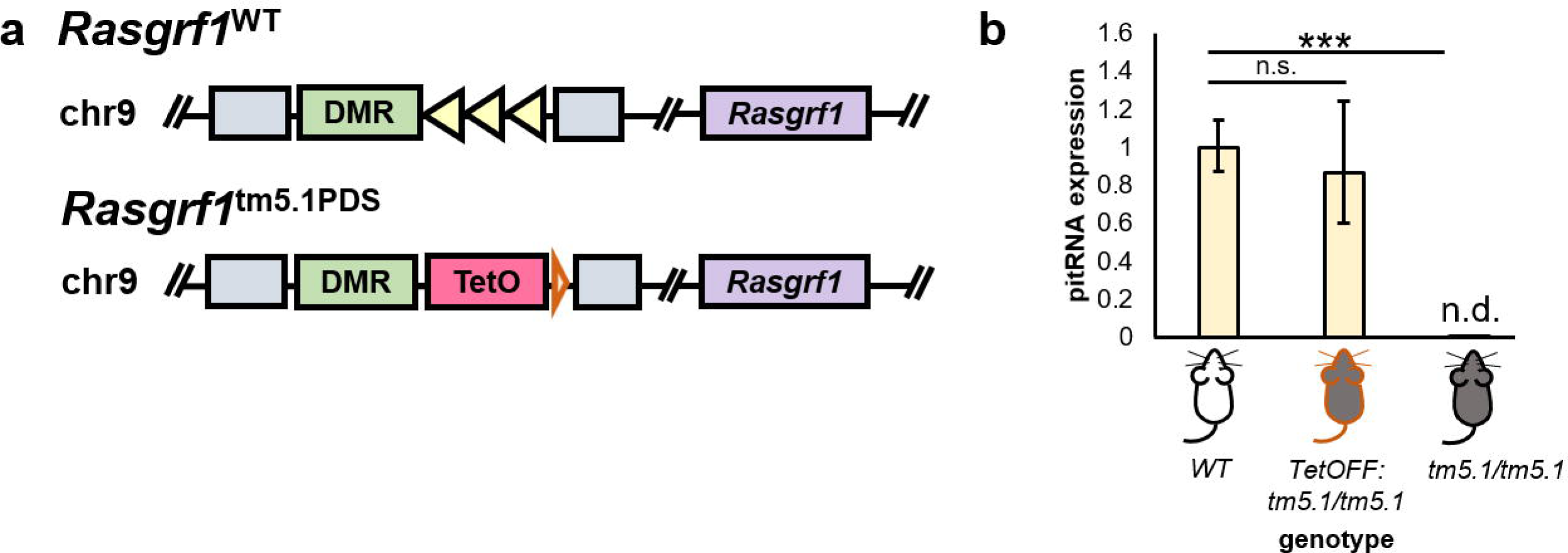
Schematics and pitRNA expression of the wild-type *Rasgrf1* and *Rasgrf1*^*tm5.1PDS*^ alleles. **a)** Schematics of the wild-type *Rasgrf1* and *Rasgrf1*^tm5.1PDS^ alleles. Green box represents the *Rasgrf1* differentially methylated region (DMR). The Imprinting Control Region (ICR) lies 30 kb upstream of *Rasgrf1* coding sequence (purple box), and includes the DMR and the 1.6kbp *Rasgrf1* repeats (yellow triangles). Grey blocks denote 5’ and 3’ homologous arms of the targeting vector pETC6, described in Methods. Orange triangle downstream of *Rasgrf1*^tm5.1PDS^ is a residual frt site. Figures are not drawn to scale. **b)** pitRNA expression postnatal day 1 testes of wild-type, *tm5.1/tm5.1*, and *TetOFF*:*tm5.1/tm5.1* animals. *TetOFF* transactivates *tm5.1* to physiological levels; *tm5.1* homozygotes express undetectable levels of pitRNA in the absence of *TetOFF*. Error bars represent standard error across biological triplicates. ***, p < 10-6; n.s., not significant; n.d., not detected at 40 cycles.

### The *tm5.1* allele lacks DMR methylation and *Rasgrf1* expression, like the *Rasgrf1*^*tm1*^ repeat-deficient allele

We first characterized *tm5.1* in the absence of a transactivator. We expected that, in the absence of a transactivating protein, *tm5.1* would neither accrue methylation at its DMR or impart imprinted expression at *Rasgrf1*, similar to the repeat-deficient allele, *Rasgrf1*^*tm1*^ (*tm1*). The *tm5.1* DMR was hypomethylated when paternally transmitted (+/*tm5.1*, Fig S2e), leading to minimal expression of *Rasgrf1* in the brain as measured by qRT-PCR (FigS2d); +/*tm5.1* animals were on average of lower body weight than wild-type littermates (Fig S2f), consistent with findings that repeat-deficient animals are underweight [41].

Also consistent with *tm1*, a portion of +/*tm5.1* animals were methylated at, and expressed *Rasgrf1* at wild-type levels from the *tm5.1* allele. Also as seen with the *tm1* allele, a portion of mice with the +/*tm5.1* genotype had *tm5.1* methylation and expression in the N2 and N3 generations (4 out of 14 animals in the N2 generation; 2 out of 15 animals in the N3 generation). Consistent with findings using repeat-deficient animals [41], this was a stochastic event, and the methylation status of *+/tm5.1* offspring was not dependent on the methylation state of their +/*tm5.1* fathers (Fig S2a-b).

### Induction of pitRNA from *Rasgrf1*^tm5.1PDS^ via the *TetON* and *TetOFF* systems

To ascertain transactivator-dependent induction of pitRNA, we then generated *tm5.1* mice expressing one of two transactivating proteins, *TetON* and *TetOFF*. As discussed in Methods, the *TetOFF* allele was generated from the commercially available *pA-TetOFF* allele, where *TetOFF* is preceded by a floxed neomycin-resistance polyadenylation cassette. The polyadenylation cassette was removed by embryonic day 6.5 by crossing pA-*TetOFF* males with females carrying a *Cre* transgene driven by the *Sox2* promoter [42]. Successful Cre-mediated recombination was confirmed by Sanger sequencing of PCR products spanning the pA-cassette (Fig S3). We expected pitRNA transcription from the *tm5.1* allele would depend either on the *TetOFF* transgene in the absence of tetracycline, or the *TetON* transgene in the presence of doxycycline. We assayed pitRNA induction in several adult tissues by endpoint PCR, as well as by qPCR in the neonatal male germline, adult liver, neonatal brain, and oocytes. pitRNA induction from the *tm5.1* allele required a transactivating protein, and was expressed at physiological levels in the neonatal male embryonic germline of males (Fig 1b), and from 10 to 1000-fold wild-type levels in adult tissues (Fig S4, Fig S7a) depending on the tissue assayed. In all tissues, pitRNA was silent in the absence of a transactivator.

### pitRNA induction in the male germline is insufficient for establishment of germline methylation at *Rasgrf1*

Having confirmed that pitRNA could be induced from *tm5.1* using both the *TetON* and the *TetOFF* systems, that expression in the neonatal germline was at physiologic levels, and that *tm5.1* was transcriptionally silent in the absence of a transactivator, we tested whether artificially regulated pitRNA expression was sufficient to impart methylation at the *Rasgrf1* DMR in the male germline, independently of the repeats that normally effect this regulation. To perform this analysis, we used gonads from male embryos heterozygous for the *tm5.1* allele that also carried the *TetOFF* or the *TetON* transgenes. We assayed methylation of the *tm5.1* and WT DMRs under conditions where the pitRNA is induced from the *tm5.1* allele. The use of heterozygotes enabled us to monitor methylation of both the *tm5.1* allele, with artificial regulation of the pitRNA from the Tet Operator, and the wild-type allele, as an internal control, which has natural regulation of the pitRNA from the repeats. gDNA was prepared from the gonocyte and somatic cell fractions of developing male gonads, and assayed for DMR methylation by targeted bisulfite sequencing (Fig 2a-d, S5a,b lower panels) and COBRA (Fig S5a,b upper panels), using assays specific for the *tm5.1* and wild-type alleles. Both assays revealed that the *tm5.1* DMR was hypomethylated in gonocytes from each of six mice tested, despite the proper regulation of pitRNA in the germline from the *tm5.1* allele. In contrast, the wild-type DMR from the same animals was hypermethylated, as expected. This pattern was observed in male gonocytes regardless of the parental modes of inheritance of the two alleles, or whether the *TetON* or *TetOFF* regulator was used. As a control for purity of germ cells, we performed BS-PCR and sequencing for the *Igf2r* DMR, which is methylated only upon maternal transmission, and found extensive hypomethylation, as expected for male germline cells [43]. We also assayed methylation states of the two alleles in somatic fractions of developing gonads. As with gonocytes, the *tm5.1* allele was unmethylated regardless of mode of inheritance or transactivator.

**FIGURE 2.**
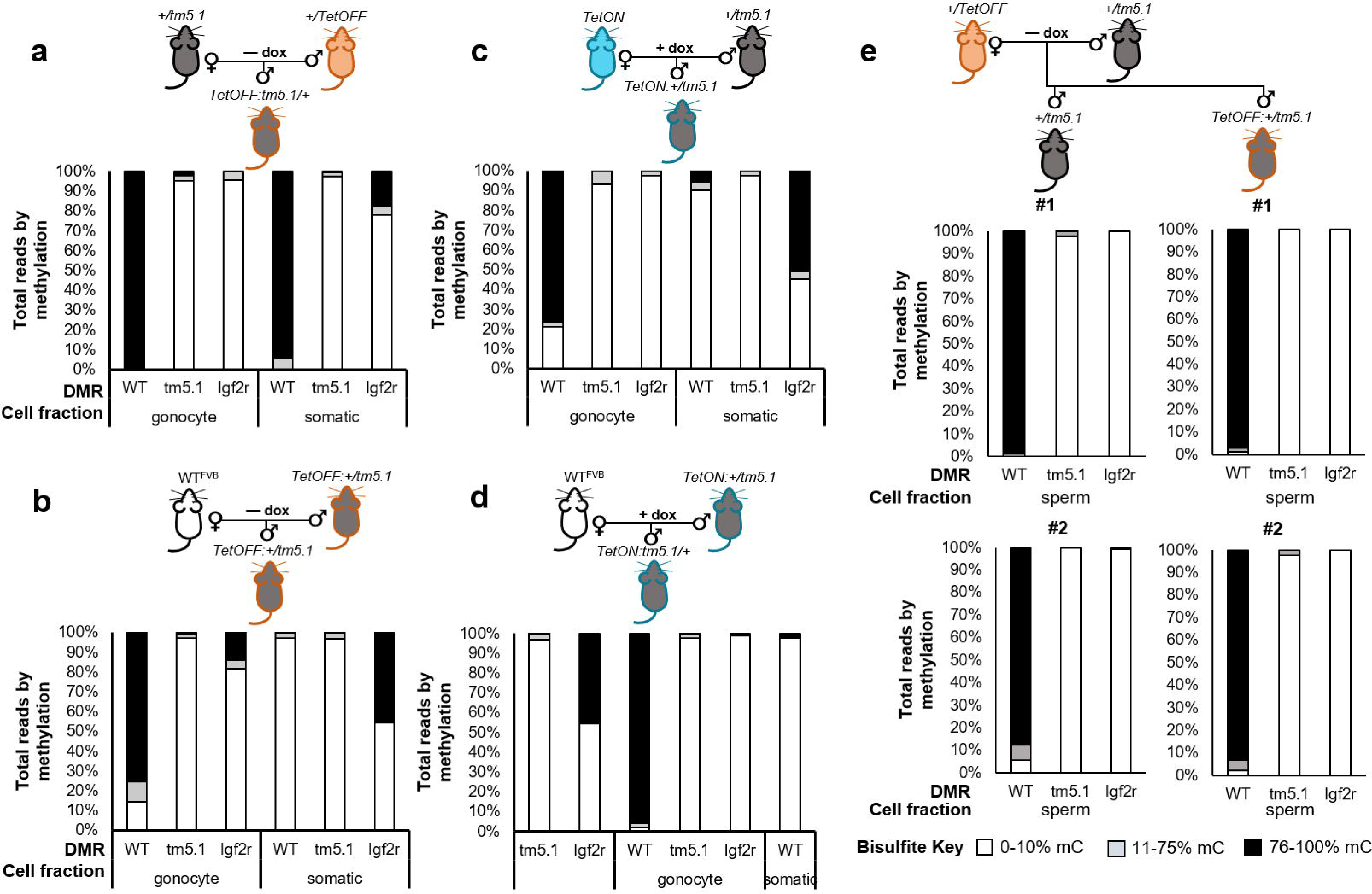
Induction of pitRNA in the male germline does not impart methylation *in cis*, at the *tm5.1* DMR, or in *trans*, at the *Rasgrf1* DMR. DNAs were collected from gonocyte and somatic fractions of male embryos (a-d) and mature sperm from adult males (e). The somatic and germline fractions of one male is shown in each panel a-d; panel e shows bisulfite analysis of the sperm of two males (#1 and #2). Animals were on the C57BL/6 or FVB/n (FVB, c, d) backgrounds. Mothers were fed chow containing (+ dox), or lacking (– dox) doxycycline between mating and birth. Bisulfite PCR was done using PCR primers specific to the *tm5.1* DMR or, as controls, the DMRs from wildtype (WT) *Rasgrf1* and *Igf2r*. WT and *Igf2r* are respectively expected to be hyper- and hypomethylated in male gonocytes; *Igf2r* is expected to be 50% methylated in soma. Number of CpG dinucleotides assayed from *tm5.1*, WT, *tm5.1*and *Igf2r* totaled 15, 17, and 12 respectively. MiSeq libraries prepared from PCR products were sequenced to a minimum of 12 reads per sample for the two *Rasgrf1* alleles (range: 12-28,067; median: 135). Data are reported as the percentage of reads showing the methylation levels indicated in the Bisulfite Key at lower right. *Igf2r* hypomethylation in the gonocyte fractions of each sample indicate minimal somatic contamination. Consistent hypomethylation of the *tm5.1* allele in each pedigree and sample indicates pitRNA expression in the absence of the repeats is insufficient for DMR methylation.

As expected, the wild-type allele was hypermethylated upon paternal transmission, and hypomethylated upon maternal transmission. Hypomethylation of the maternal DMR in somatic fractions demonstrated that pitRNA expression from the *tm5.1* allele does not act in *trans*.

We further assayed *tm5.1* and wild-type DMR methylation in mature sperm of *tm5.1* heterozygotes, where the pitRNA was regulated by *TetOFF* in the absence of doxycycline. As with gonocytes, the *tm5.1* DMR was consistently hypomethylated, whereas the wild-type DMR was methylated (Fig 2e and Fig S5c). We concluded that in the embryonic and mature male germlines, pitRNA expression alone was not sufficient to impart methylation at the *tm5.1* DMR in *cis*, indicating that the repeats perform an additional necessary function for DMR methylation, beyond controlling pitRNA expression.

### pitRNA induction in the male germline is insufficient for somatic methylation at *Rasgrf1*

In previous studies, exporting the *Rasgrf1* ICR to the *Wnt1* locus led to hypomethylation of the mutant allele in sperm, but hypermethylation in somatic tissue after fertilization [37]. Though sperm methylation at the modified *Wnt1* allele was higher than sperm methylation at the *tm5.1* allele, we determined if expression of pitRNA by TetO induction of the *tm5.1* allele could enable methylation in somatic tail DNA of progeny after paternal transmission. In all tail samples tested, the *tm5.1* allele remained unmethylated regardless of which transactivator was used to control pitRNA (Fig 3a-b). These findings demonstrated that like methylation in the male germline, methylation in somatic tissue of offspring after paternal transmission is not enabled by pitRNA expression alone. Instead, and consistent with findings from the *Wnt1* mutant allele, additional features of the repeats, beyond their control of pitRNA expression, are necessary for methylation.

**FIGURE 3.**
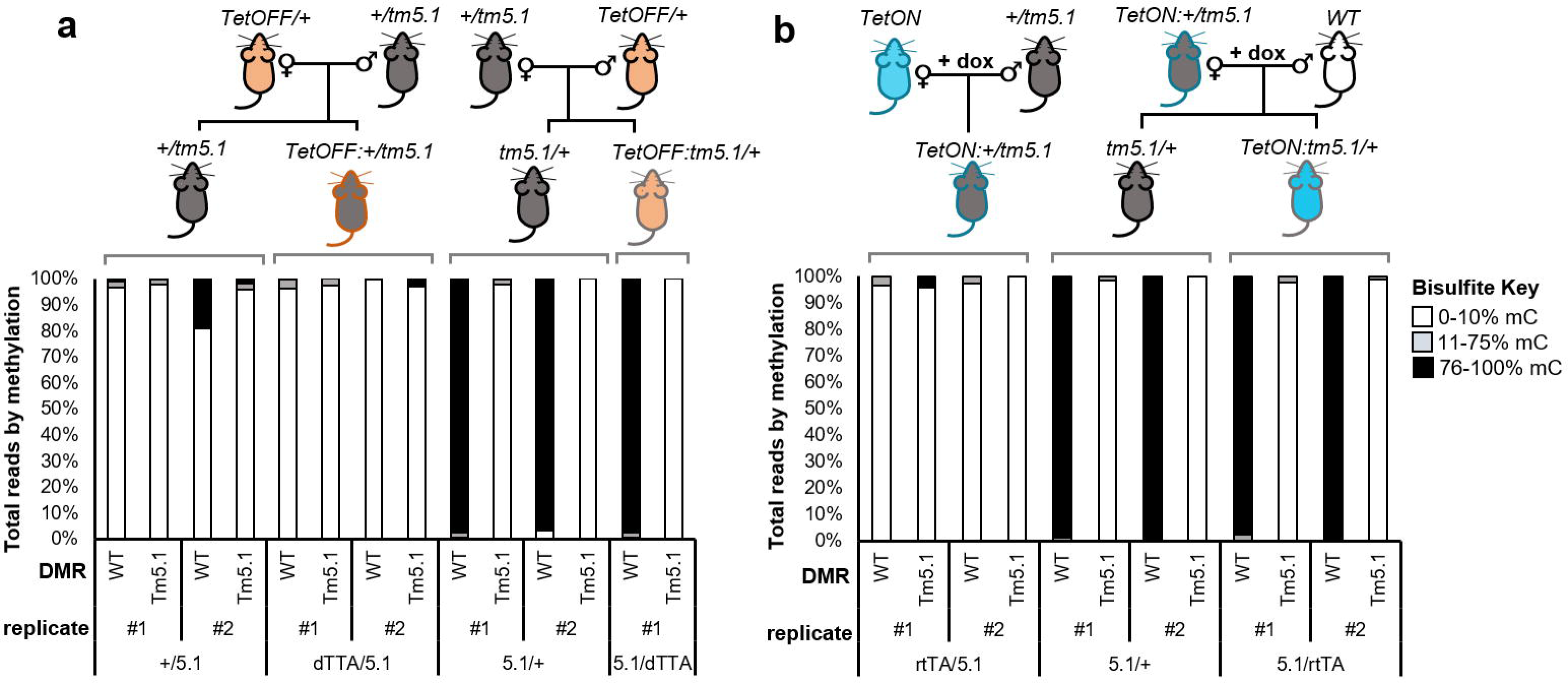
*TetOFF*-mediated transactivation does not affect *tm5.1* DMR methylation in neonatal tail. **a**) Targeted bisulfite analysis of WT and *tm5.1* DMRs in neonatal tail gDNA; pedigree shown at top. *tm5.1* is hypomethylated regardless of parental descent and presence of *TetOFF*. The WT DMR is methylated depending on parent of origin—if inherited maternally, the wildtype DMR is hypomethylated; if inherited paternally, the wildtype DMR is hypermethylated. Two biological replicates for all genotypes are shown (#1 and #2) except for the *TetOFF:tm5.1/+* genotype, where one animal is shown. **b)** Targeted bisulfite analysis of 5.1/+, *tm5.1*/*TetON*, and *TetON*/*tm5.1* neonatal tail gDNA. In all genotypes, *tm5.1* DMR is hypomethylated; as expected, the wild-type DMR is methylated if inherited paternally as in *tm5.1* and *tm5.1*/*TetON* animals and hypomethylated if inherited maternally as in *TetON*/*tm5.1* animals. Reads for two animals (#1 and #2) per genotype are shown. **, p < 10^−3^; ***, p < 10^−6^; n.s., not significant.

### *tm5.1* transactivation and pitRNA induction leads to expression changes of neighboring genes

We expanded our initial analysis of methylation by characterizing expression states of *Rasgrf1* and nearby loci in mice carrying the *tm5.1* allele. *Rasgrf1* expression in neonatal brain requires either methylation of the DMR, which is a methylation-sensitive enhancer blocker, or ectopic insertion of an enhancer proximal to the *Rasgrf1* promoter [44]. We found that *TetOFF*-mediated *tm5.1* induction led to a tenfold upregulation of *Rasgrf1* in neonatal brain relative to wild-type regardless of the parental origin of the *tm5.1* allele. Sequencing *Rasgrf1* RT-PCR products revealed that *TetOFF*:+/*tm5.1* animals that inherited *tm5.1* paternally, expressed *Rasgrf1* solely from the paternal *tm5.1* allele (Fig 4a-c). *TetOFF: tm5.1*/+ animals that inherited *tm5.1* maternally expressed *Rasgrf1* from both the maternal and paternal alleles at a ratio of approximately 9:1, indicating there was a dramatic upregulation of *Rasgrf1* from the normally silent wildtype maternal allele, when it was replaced by *tm5.1*, and with transactivation by *TetOFF*. This was accompanied by continued *Rasgrf1* expression from the paternal wild-type allele (Fig 4a-c). We observed similar effects with *TetON* transactivator. Though the magnitude of *Rasgrf1* induction was lower, *TetON* also activated expression from the maternal *tm5.1* allele (Fig 4 d-f, Fig S6).

**FIGURE 4.**
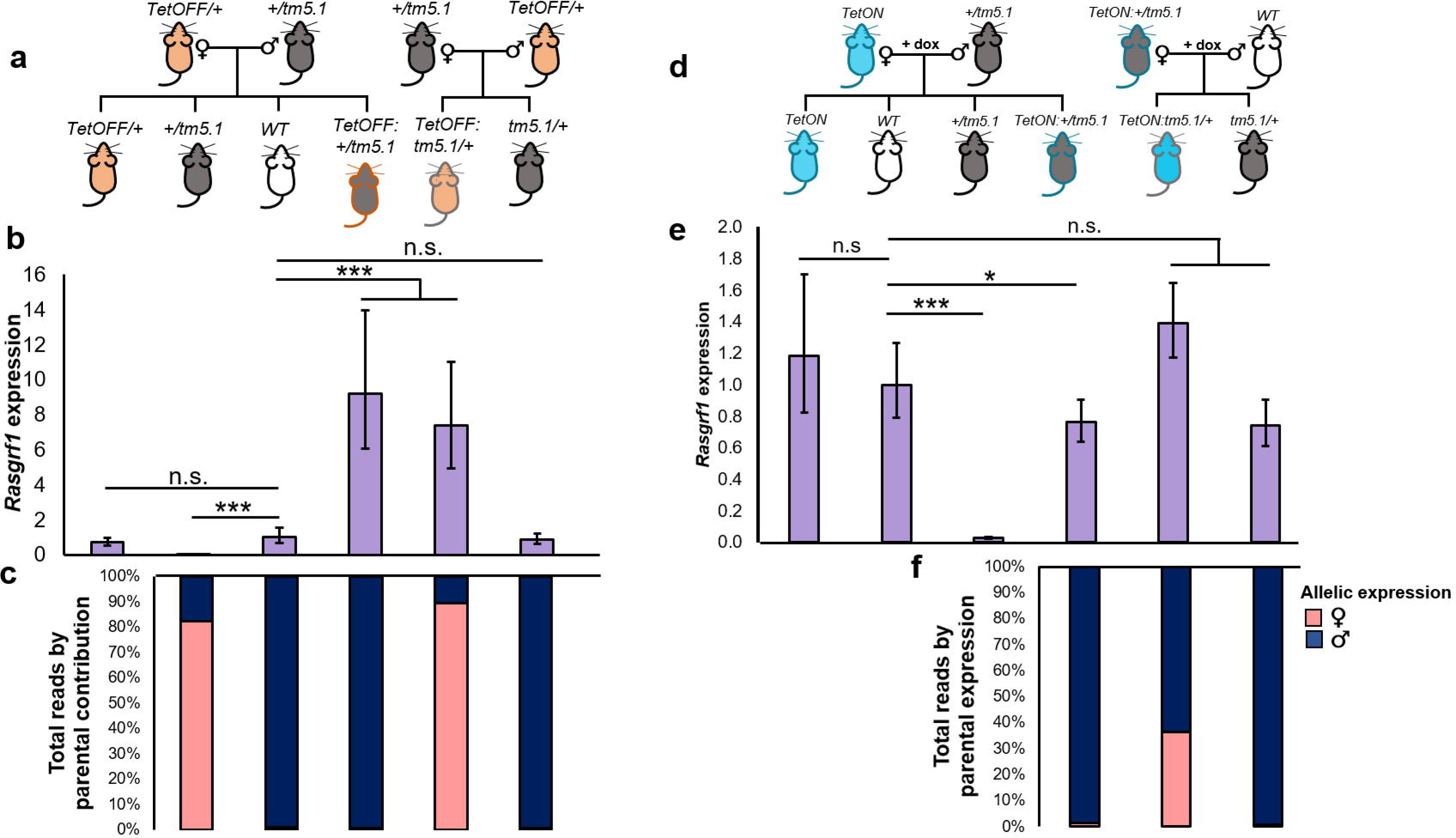
Transactivation of *tm5.1* upregulates *Rasgrf1* expression from *tm5.1* in neonatal brain. **a** and **d)** Pedigrees of animals used respectively for **b**, **c)**, and **e**, **f). b, e)** *Rasgrf1* expression levels were assayed by qRT-PCR using neonatal brains from the pedigrees, and genotypes shown directly above in **a** and **d)**. Error bars represent standard error from, at minimum, biological triplicates. *Rasgrf1* levels were normalized to *Rpl32*, and levels in WT mice were arbitrarily set at 1. **c** and **f)** Relative expression of *Rasgrf1* from the two parental alleles (maternal pink; paternal, blue). Allelic assignments were made by sequencing libraries of RT-PCR products amplified using primers that span *Rasgrf1* polymorphisms, which distinguish *tm5.1*, made on the 129S4 background, and the WT allele, contributed by *TetOFF* or *TetON* mice on the C57Bl/6 background. Products were sequenced to a minimum of 15 reads per animal from at least two biological replicates. *Rasgrf1* is dramatically downregulated in *+/tm5.1* animals, that inherit *tm5.1* paternally, with most residual expression coming from the largely silent maternal allele. The *tm5.1* allele increases *Rasgrf1* expression in a *Tet* transactivator-dependent manner, regardless of the parental origin of *tm5.1*. No doxycycline treatments were applied for **c** and **d)**, but were applied for **e** and **f)**. *, p < 0.05; **, p < 0.01; ***, p < -6 10.

The *Rasgrf1* locus lies within two overlapping annotated regions of chromatin interaction, a smaller 150 kb and encompassing 250 kb region, as shown by cohesin ChIA-PET analysis of mouse embryonic stem cells [45] (Fig 5a). To define the extent of *TetOFF*-mediated transactivation, and its relationship to the bounds of known regions of interaction, we queried the effects of *TetOFF* on other transcripts within the interacting regions. In neonatal brain, all transcripts within the minimal 150kb interaction domain were upregulated in both *TetOFF*:+/*tm5.1* and *TetOFF*:*tm5.1*/+ animals, in a transactivator dependent manner, indicating the effects of transactivation extended throughout the 150kb domain. Interestingly, *TetOFF*-dependent effects on brain expression of transcripts within the *tm5.1* domain varied, depending on parental origin of *tm5.1*, consistent with the existence of distinct chromosomal architecture for parental alleles within imprinted regions (Fig 5b). The allele-specific effects were even more dramatic when expression was assayed in neonatal testes: transcripts linked to *tm5.1* that were upregulated by *TetOFF* when *tm5.1* was maternally transmitted were downregulated when it was paternally transmitted. Additionally, the effects extended over a broader chromosomal domain in testes, and on the paternal chromosome, highlighting parental- and tissue-specific chromatin architecture at *Rasgrf1* (Fig 5c).

**FIGURE 5.**
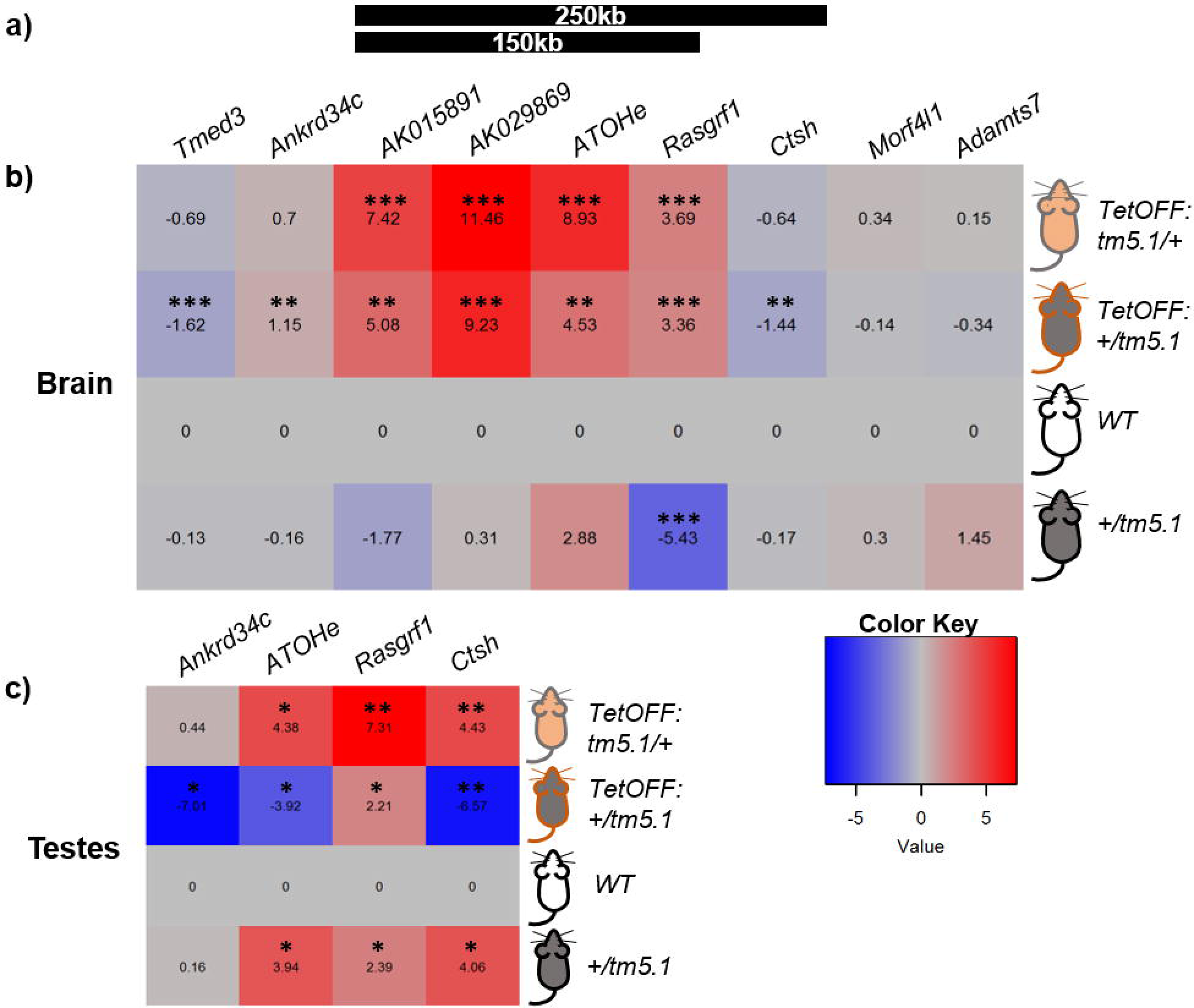
Regional transcription is perturbed in *TetOFF*: +/5.1 and *TetOFF*: 5.1/+ brain and testes. **a)** Two regions of chromatin interactions (black bars) are annotated at the *Rasgrf1* locus as predicted by cathepsin ChiA-PET in mouse embryonic stem cells [45]; locations are shown relative to the genes indicated below in **b)**. Lengths of each predicted interaction are shown in kilobases (kb) within each bar. Relative log 2 expression levels in neonatal brain **b)**, and neonatal testes **c)** of transcripts from chr9:89,60,000-90,100,000 are shown, displayed left to right as they are located 5’ to 3’. In brain, *TetOFF* exerts dramatic upregulation of *Rasgrf1* and nearby transcripts *AK015891*, *AK029869*, and an annotated ATOH1 binding site 3’ of the *Rasgrf1* repeats (*ATOHe*) in mice with both paternally and maternally inherited *tm5.1*. However, distant transcripts *Tmed3* and *Ctsh* are downregulated modestly in brains of offspring inheriting *tm5.1* maternally. **c)** Log10 expression of four transcripts (a subset of the nine assayed in brain) in neonatal testes. While *ATOHe* and *Ctsh* are upregulated when *tm5.1* is transactivated and paternally inherited, *Ankrd34c*, *ATOHe*, and *Ctsh* are downregulated when *tm5.1* is transactivated and inherited maternally. All expression data are relative to -6 Rpl32. **Color Key** at lower right. *, p < 0.05; **, p < 0.01; ***, p < 10.

### pitRNA loading of oocytes does not produce paramutation

Previously, our lab described a paramutation-like phenomenon at *Rasgrf1* associated with the *Rasgrf1*^*tm3.1PDS*^ allele (*tm3.1*), in which the repeats were replaced by the imprinting control region (ICR) of *Igf2r* [43]. Progeny carrying a paternal *tm3.1* allele exhibited a derepression of the maternal allele, as interpreted from endpoint RT-PCR; and further, some wild-type offspring of +/*tm3.1* females demonstrated continued depression of the maternal allele, though they lacked the original paternal allele that incited the derepression. This intergenerational effect is a key feature of paramutation. The *Igf2r* sequences in the *tm3.1* allele harbors the promoter for *Air*, a non-coding RNA that regulates imprinted expression at the locus [46]. Its presence and orientation in the *tm3.1* allele could impart novel expression patterns of the pitRNA; accordingly, we hypothesized that the intergenerational, paramutation-like effects could involve oocyte loading of pitRNA. To test this hypothesis, we treated females carrying the *tm5.1* allele, and the TetON activator, with intraperitoneal dox for three days, which led to oocyte pitRNA levels approximately 90-fold higher than in wild-type mice (Fig S7a). Females subjected to these treatments were bred to wild-type males, and maintained on dox chow for the duration of pregnancy—as such, their *TetON:*+/*tm5.1* offspring were informative for somatic effects of pitRNA induction in oocytes (Fig S6). Additionally, their wild-type, *TetON:*+/+, and +/*tm5.1* offspring were informative for determining if pitRNA loading in oocytes could induce paramutation. In none of these offspring born to mothers with pitRNA preloaded in their oocytes did we observe effects on imprinting status or expression levels of *Rasgrf1* (FigS7c, d). COBRA analysis of the maternal *tm5.1* allele transmitted by oocytes preloaded with pitRNA showed that it remained hypomethylated (Fig S7e). We concluded that preloading oocytes to nearly 100-fold levels of pitRNA relative to wild-type was insufficient to induce transgenerational effects.

## DISCUSSION

### A model for DNA methylation at *Rasgrf1*

Despite the fundamental roles that DNA and histone modification states have for genome regulation, there have been only a handful of sequence elements identified that exert *cis*-acting control over these modification states [16-21]. We previously identified one such element at the *Rasgrf1* locus: It is a series of tandem repeats found 30 kb upstream of the imprinted *Rasgrf1* coding sequence, and lying immediately adjacent to the DMR, that are required both in the male germline for establishment of imprinted methylation at the DMR in sperm [21], and also in the pre-implantation embryo for maintenance of imprinted methylation in somatic lineages [23]. In addition to serving these functions, the repeats serve as a promoter for expression of the pitRNA in the male germline that spans the DMR, and which is processed into piRNAs. The distinct effects of lncRNAs, and the *cis*-elements that regulate them, have proven difficult to uncouple [47,48] with some success coming from the truncation of the lncRNA transcript while keeping the *cis* element intact [49,50,51,52]. To assess the activity of the pitRNA separately from the repeats, we uncoupled the two by replacing the repeats with an artificial promoter based on TetO sequences (*tm5.1*), which regulated the pitRNA to physiologic levels in the developing male germline using the *TetON* and *TetOFF* transactivators. Of nine male embryos with this pitRNA regulation analyzed, only one displayed partial gonocyte and somatic methylation of *tm5.1*. This frequency is consistent with the frequency of DMR methylation observed in mouse models lacking the repeats [23, 41]. Properly regulated pitRNA expression from the *tm5.1* allele also failed to enable methylation in mature sperm, or in somatic DNA of progeny inheriting the *tm5.1* allele from their fathers. We therefore conclude that the pitRNA is insufficient to impart DNA methylation in *cis* and that *Rasgrf1* methylation requires critical functions in the repeats, separate from pitRNA expression (Fig 6).

**FIGURE 6.**
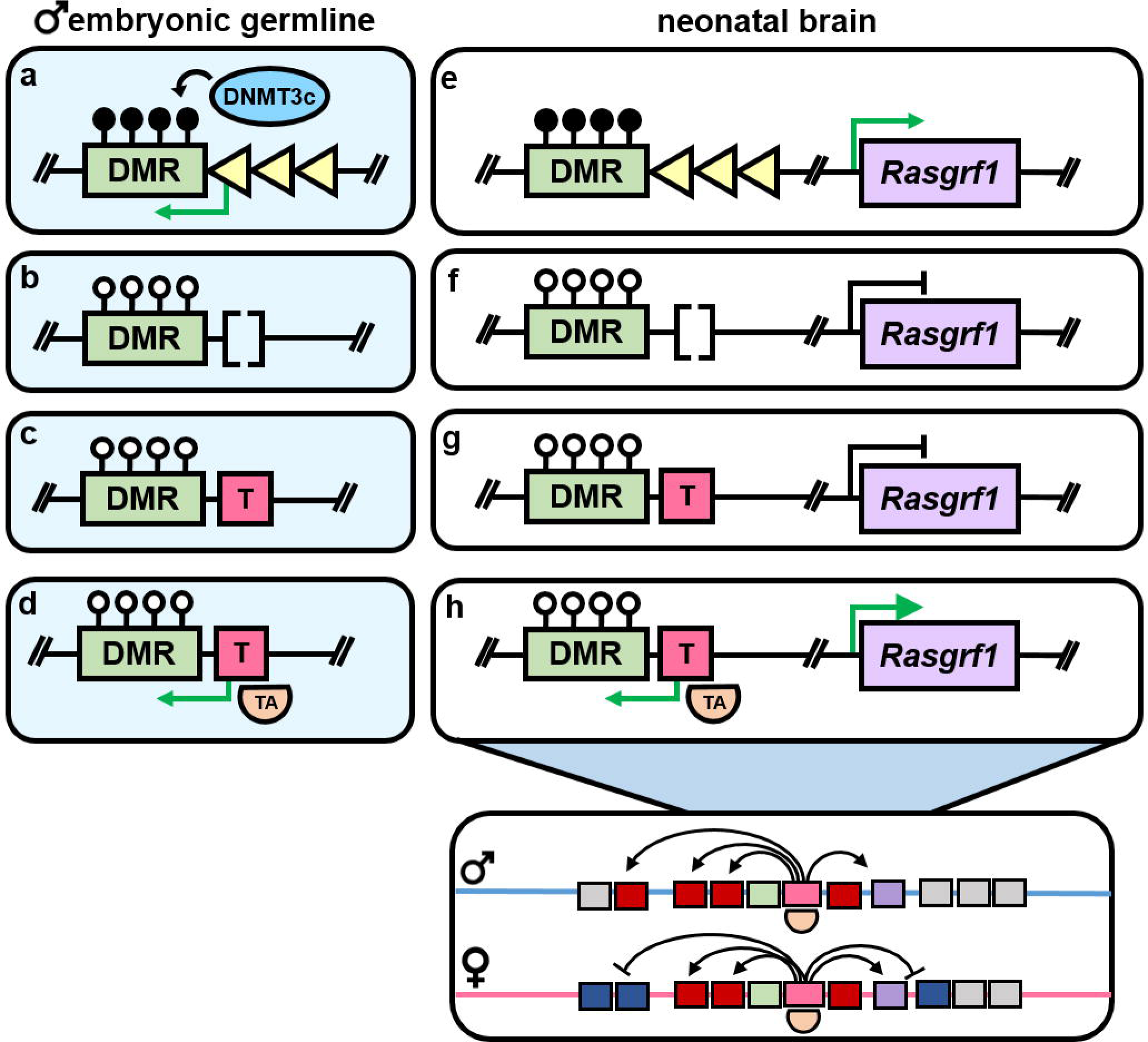
Working model for regulation of methylation at *Rasgrf1* in the male embryonic germline and neonatal brain. For neonatal brain, only the paternal allele is depicted. **a)** At the wildtype *Rasgrf1* DMR, the *Rasgrf1* repeats (yellow triangles) drive expression of the pitRNA antisense to the DMR (transcription depicted by green arrow) and increase the likelihood that the DMR is hypermethylated (filled lollipops), likely by Dnmt3c, in the male germline. **b)** *Rasgrf1*^*tm1*PDS^ (tm1) lacks the *Rasgrf1* repeats (open brackets). pitRNA is not expressed and methylation is not established at the DMR in the *tm5.1*PDS germline (open lollipops). **c)** *Rasgrf1* (*tm5.1*) is not methylated in the male *tm5.1*PDS germline. **d)** *Rasgrf1* transactivation with *TetOFF* (orange half circle labelled TA) induces pitRNA expression but does not impart methylation to the *tm5.1* DMR. **e)** In neonatal brain, *Rasgrf1* is paternally methylated and expressed. **f, g)** *tm1* and *tm5.1* brains have severely decreased *Rasgrf1* expression. **h)** transactivation of tm5.1 causes strong upregulation of *Rasgrf1* but the *tm5.1* DMR remains hypomethylated. **Outset:** *TetOFF*-mediated transactivation of *tm5.1* affects nearby transcription differentially depending on parental inheritance (blue and pink lines). Upregulated and downregulated genes are depicted by red and blue boxes respectively; unchanged genes are depicted in gray. TetO and DMR are indicated by the pink and green boxes respectively; *TetOFF* binding is indicated by the orange half circle.

Our findings that pitRNA expression is insufficient to control imprinted methylation of *Rasgrf1* are consistent with other work from our lab, in which the *Rasgrf1* DMR and repeats were exported to an ectopic site at the *Wnt1* locus. Paternal transmission of the modified *Wnt1* allele enabled complete methylation of the ectopic DMR in somatic DNA of the offspring despite germline expression of the pitRNA at only 1% of the levels seen from the endogenous *Rasgrf1* locus; methylation of the *Wnt1* DMR in sperm DNA at only 30% of the levels found at the *Rasgrf1* DMR.

One interpretation of the present study, combined with the findings from the *Wnt1* study is that the pitRNA is irrelevant to *Rasgrf1* methylation: that instead, the repeats are both necessary and sufficient for this function, and the transcription of the pitRNA is an irrelevant sequela to some inherent activity at the repeats [reviewed in 53]. However, other data from our lab support a role for the pitRNA [24], likely by influencing the efficiency and/or timing of *Rasgrf1* methylation in a probabilistic way rather than in a deterministic way. This characterization is similar to the *Xite* element at the X-Inactivation Center, which is transcribed, and mediates the probability with which an X chromosome undergoes X chromosome inactivation [54]. The probabilistic model for pitRNA function is also consistent with our findings that mice deficient for piRNA binding proteins MILI and MIWI2, as well as the piRNA biogenesis factor MitoPLD, have diminished methylation at the *Rasgrf1* DMR, though methylation is not completely lost [24]. Unless the piRNA factors are exerting their effects through the *Rasgrf1* repeats, and not through the pitRNA they in fact target for piRNA processing, the pitRNA is very likely to contribute to DMR methylation, despite not being absolutely required.

We also utilized the *tm5.1* allele to query whether previously observed intergenerational effects could be due to oocyte preloading of the pitRNA. Others have reported that gametic loading of RNAs could impart such effects to the next generation [35,36]. We achieved oocyte pitRNA levels up to 90-fold greater than wild-type by intraperitoneal doxycycline administration, however, this produced no effects on imprinted *Rasgrf1* expression or methylation in the wild-type offspring of females subjected to oocyte preloading of pitRNA.

The repeats have several features that might enable their control of local methylation states, separate from their control of pitRNA expression. Besides being highly repetitive, the GC-richness of the repeats might be sufficient to recruit methylation [55,56] which then spreads into the DMR. This is seen at *H19* where somatic methylation on the paternal arises adjacent to the germline DMR [57], and occurs when CTCF is unbound to the germline DMR [58,59,60,61,20). Within the repeats are two canonical binding sites for the transcription factor Sp1, which besides its known role in the regulation of gene expression [62], can mediate chromatin structure through the recruitment of chromatin remodeling factors [63,64] and mediating enhancer-promoter interactions [65,66]. Sp1 is known to bind the secondary DNA structure G-quadruplexes (G4s) [67,68], which the *Rasgrf1* repeats are predicted to form, as well as its canonical sequence. Recently, a G4 was characterized at the imprinted *H19* locus. Binding of Sp1, in conjunction with the G4, suppressed *H19* transcription [69]. To our knowledge, G-quadruplex formation at other imprinted loci beyond *H19* has not been investigated.

However, differential G4 formation in the maternal and paternal germlines could be a platform upon which the distinct chromatin states observed at the maternal and paternal DMRs are built.

Our lab previously showed that, at the *Rasgrf1* DMR, H3K27Me and DNA methylation are mutually antagonistic, whereby the presence of one mark blocks the deposition of the other [22]. Placement of H3K27me3 requires YY1 and PRC2, which, in *Drosophila* [70] and likely in mammals [71], can be recruited to Sp1 sites. It is possible in the male germline, that Sp1 binding to the repeats excludes YYI and PRC2 binding, enabling paternal allele methylation.

Other factors required for parent-of-origin specific DNA methylation might require the repeats for their recruitment. The GHKL ATPase Morc1 is a known repressor of transposable elements in the embryonic male germline; *Morc1*-null mice are hypomethylated at the *Rasgrf1* DMR [72]. The KRAB-domain containing zinc-finger binding protein ZFP57 and its cofactor Trim28 [73] are typically thought of as an imprinting maintenance mechanism. Loss of zygotic Trim28 disrupts imprinting at *Rasgrf1* along with many other maternally and paternally imprinted loci [74].

Interestingly, KRAB-domain ZFP binding can also trigger *de novo* DNA methylation during mouse embryogenesis [75].

### Additional technical considerations for the study of imprinted loci, and use of transactivators

Our experiments highlight an important technical consideration for the imprinting field, that allele-specific expression analyses must be supported with quantitative data.

By end-point RT-PCR followed by allele-specific restriction digest, *TetOFF*-mediated transactivation of maternally inherited *tm5.1* appears to fully reverse imprinting, with maternal-only bands upon gel electrophoresis (Figure S7). Similar data have previously been interpreted as a silencing of the paternal allele *in trans* [76]. However, sequencing of *Rasgrf1* cDNA of *tm5.1/+:TetOFF* neonatal brain demonstrated that roughly 10% of total *Rasgrf1* reads are paternal in origin, suggesting that the wild-type paternal allele continued to express *Rasgrf1* at wild-type levels, and was not silenced *in trans*.

Similarly, end-point analysis of *+/tm5.1* animals lacking a transactivating allele revealed biallelic expression. While this could be interpreted as a possible *trans* effect of the paternally inherited *tm5.1* allele exerting effects on the maternally inherited and normally transcriptionally silent WT allele, qPCR of *Rasgrf1* in these animals demonstrated severely reduced *Rasgrf1*, suggesting instead that, in the *tm5.1* system, biallelic expression reflects minimal transcription.

An unexpected outcome of this work was that *TetOFF*-mediated transactivation of *tm5.1* influenced expression across the entire chromatin domain, defined by cohesin ChiA-PET [45], where the transactivator bound, and that this influence was both allele- and tissue-specific. The cohesin ChiA-PET study defined two regions of interaction, which may represent the maternal and paternal alleles. These widespread transcriptional changes were not observed in *TetON:+/tm5.1* tissues (data not shown), suggesting that these widespread effects may be restricted to the *TetOFF* system. While this finding was not an objective of our study, it emphasizes the considerations that should be taken when interpreting transcriptional effects using potent transactivating systems such as the *TetOFF* protein.

In conclusion, our data support the existence of a second, pitRNA-independent mechanism for DNA methylation at *Rasgrf1*. We propose a *cis*-acting mechanism by which the repeat sequences themselves are largely responsible for methylation control at the locus, and the pitRNA increase the probability of methylation in the paternal germline.

## MATERIALS AND METHODS

### Primer sequences for all analyses are listed in Table S1

#### TetO vector generation

We modified pYP1, which carries the DMR and *Rasgrf1* repeats, and 4 kb of homologous flanking sequence [77] to carry seven copies of the Tet Responsive Element (collectively termed TetO) in place of the *Rasgrf1* repeats as follows. The *Rasgrf1* repeats were removed *via* restriction digest with *Cla*I and *Mlu*I; sticky ends were blunted with Klenow; the plasmid backbone was gel purified and then ligated closed generating pDHT2. pDHT2 and pPX3, a vector containing the Tet Operator, were digested with *Nhe*I, and linearized pDHT2 was ligated to the TetO sequences, generating pDHT3, which was confirmed by Sanger sequencing. The 3’ homologous arm of pDHT3 was then shortened to approximately 1kb by restriction digest with *Bsr*GI and *Sfi*I to generate pETC6, which was linearized with *Pci*I prior to lipofection into ES cells.

### CRISPR/Cas9-mediated generation of TetO^tg^, TetOIZI ^tg^, *Rasgrf1*^tm5.0PDS^ and *Rasgrf1*^tm5.1PDS^

pX330 (Addgene Plasmid # 42230) was modified to carry PDS 2195-6, a complementary primer pair coding for an sgRNA targeting the *Rasgrf1* repeats, following the Zhang lab protocol [78] to generate pX330-rep5. PDS 2195-6 was designed using the CRISPR Design Tool (http://crispr.mit.edu:8079/). To effect homology directed repair at the *Rasgrf1* locus, v6.5 embryonic stem cells [79] were lipofected with linearized pETC6 and pX330-rep5 with Lipofectamine 2000, following the manufacturer’s protocol. Cells were allowed to recover overnight, then treated for 10 days with 300 ug/mL G418 (Sigma A1720). Colonies were picked and genotyped with PCR 2359-8 and PDS 2344-2263, which generate a product only from the targeted allele (*Rasgrf1*^tm5.0PDS^) and PDS 2757-8, an internal PCR for TetO, to detect cells harboring a randomly inserted vector that provided the transgenic (Tg) model. Targeted and transgenic status for tm5.0 and TetO^Tg^ were further confirmed by Southern blot [21].

TetO^Tg^ and *Rasgrf1*^tm5.0PDS^ ES cells were microinjected into B6(Cg)^Tyrc-2J^/J blastocysts by the Cornell University Transgenics Core. 22 chimeras were recovered. Germline transmission was confirmed by diagnostic crosses to FVB/N females and PCR with PDS 2757-8. Chimeras were crossed with mice constitutively expressing FlpE recombinase (JAX Strain 003800) to generate *Rasgrf1*^tm5.1PDS^ and TetO⍰ ^Tg^.

Recombination was confirmed by Sanger sequencing of PDS 2262-3 PCR products, which spans the neo resistance cassette and frt sites.

### Generation of *TetOFF* mice

To produce mice that constitutively express the tet transactivator (*TetOFF*), mice carrying a tTA transgene preceded by a floxed neomycin-polyadenylation cassette (pA-*TetOFF*, JAX Strain No 011008) were crossed with mice constitutively expressing *Sox2*-Cre (JAX Strain No 008454). Recombination and subsequent loss of the neomycin resistance cassette was confirmed by endpoint PCR with PDS 2794-5, followed by Sanger sequencing. PDS 2794-5 sequences were supplied by Bruce Morgan [80].

### Induction of pitRNA

#### TetOFF

*tm5.1* mice were crossed to *TetOFF* mice.

Induction of pitRNA expression was validated in adult and neonatal tissues using PDS 2266-7.

#### TetON

Female mice were injected with 0.01 mg/g body weight of 0.01 mg/mL doxycycline hyclate (Sigma D9891) intraperitoneally every 24 hours for three days as a preloading phase, then bred. Breeding pairs were fed 200 mg/kg doxycycline chow (BioServ S3888) for the duration of pregnancy. Induction of pitRNA expression was validated in all tissues by endpoint and qRT-PCR using PDS 2266-7 and PDS 2916-7, a primer pair specific for pitRNA produced from the *tm5.1* allele.

### Allele-specific analysis of *Rasgrf1* expression

Neonatal brains were collected at postnatal day 2; the olfactory bulbs were visually identified under dissection microscope and discarded. Other tissues were collected by gross dissection. Tissues were snap frozen in liquid nitrogen, then submerged in 1 mL Trizol. Tissues were homogenized using a Biospec Mini-Bead Beater-8 using 1 3mm steel bead in XXTuff Microvials (BioSpec XX0TX). Total RNA for all samples was processed via the Trizol protocol (Thermo Fisher Scientific 15596018). RNA was DNAse treated, random primed, and reverse transcribed using Promega RQ1 DNAse and RQ RTase (Promega M6101 and A5003 respectively) following the manufacturer’s protocol. RT-PCR was performed with PDS 245-6 (95ºC 2min, 40 cycles of 95ºC 30s, 60ºC 30s, 72ºC 30s, 72ºCC 7min using Promega GoTaq in a volume of 25 uL (Promega M3001). *For allele-specific restriction digest*: 20 uL of PCR product was digested with 2.5U *Aci*I. Digestion products were separated by electrophoresis on a 4% agarose gel. *For allele-specific read quantification*: End-point PCR samples were submitted for sequencing in MiSeq libraries as described below in. Targeted sequencing analysis. Read quantification for *Rasgrf1*-specific expression was performed on trimmed samples with the grep and wc functions. Grep sequences are listed in Table S4. MGI SNP IDs and flanking sequences are listed in Table S5.

### qRT-PCR and heat map generation

qRT-PCR was performed in 20 uL reactions using SYBR Green Master Mix (CAT 4367659) on a Biosystems 7500 with annealing temperature 60ºC for forty cycles followed by a dissociation stage. The following primer pairs were used: PDS 2266-7 for general pitRNA expression; PDS 2916-7 for TetO-pitRNA expression; PDS 2877-8 for *Rasgrf1* expression; PDS 72-3 for *Rpl32* expression; PDS 2719-8 for *ATOHe* expression; PDS 3211-2 for *Ctsh* expression; PDS 2178-9 for *Ankrd34c* expression. Heat maps were generated in R [81].

### Gonocyte collection

Females were checked for plugs, weighed to confirm pregnancy, and sacrificed at gestational day 16.5. Gonocytes from male embryos were collected as in Watanabe *et al*, 2011 [24] with some modifications. Briefly, embryonic testes were collected and incubated in 50 uL 0.25% trypsin for 10 minutes. Samples were then triturated and visually inspected for tissue disaggregation. Incubation and trituration were repeated up to two times until full disaggregation was achieved; any remaining clumps were manually removed. Samples were then transferred to McCoy’s basic medium supplemented with FBS and pre-plated for 1.5 hours. Suspended germ cells were harvested, pelleted at 300 x *g* for 8 minutes, then processed for RNA and DNA. We validated the purity of gonocytes by bisulfite sequencing of *Igf2r*, which is expected to be extensively hypomethylated in the male germline [**Error! Bookmark not defined.**].

### Oocyte collection

28 to 42 day old females were superovulated with 5 IU human chorionic gonadotropin (Millipore 367222) followed by pregnant mare serum gonadotropin (Millipore 230734) 48 hours later. Oocytes were collected the next morning via standard methods [82] and processed for total RNA via Trizol. Oocyte recovery was evaluated using RT-PCR primers for Zp3 (PDS 2212-3).

### Genomic DNA extraction from tails and cells

All DNA was collected via overnight incubation at 55ºC in Laird’s Lysis Buffer [83] and 20 ug/mL Proteinase K followed by isopropanol precipitation and resuspension in TE. Scant gDNA samples, such as those from gonocytes, were co-precipitated with 20 ug glycogen (Thermo Fisher R0551) and spun at 20,817 *x g* for 15 minutes.

### DNA extraction from sperm

The caudal epididymis and vas deferens of adult male mice were harvested and placed in PBS for 1 hr at 37ºC. Large tissue chunks were manually removed and the remaining sperm were pelleted at 400 x g for 12 minutes at 4ºC. Supernatant was discarded; the pellet was resuspended in the remaining supernatant and incubated with 1 mL somatic cell lysis buffer [84] for 1 hr on ice. Lysed samples were spun at 20,817 x g for 3 minutes at 4ºC and the supernatant was discarded. The sperm pellet was resuspended in remaining supernatant; 2 uL was mixed with Trypan Blue and examined microscopically for remaining somatic contamination.

The remaining suspension was mixed with 500 uL Buffer RLT (Qiagen 79216) and 150 mM DTT, then homogenized with 2 2mm steel beads following the protocol of Wu *et al* [85]. DNA extraction then proceeded as described above.

### Bisulfite conversion, BS-PCR, and COBRA

Bisulfite conversion was performed using the Zymo Research EZ DNA Methylation-Lightning Kit (Zymo Research D5031). BS-PCR was performed with PDS 271-272 for the wild-type DMR, PDS 272-2627 for the *tm5.1* or Tg DMR, PDS 2934-5 for the *Igf2r* DMR, and 271-287 for the *tm1* DMR using NEB Epimark HotStart *Taq* DNA Polymerase (NEB M0590) following the following cycling parameters: 95ºC for 30s, 40X (95ºC 15s, 55ºC 30s, 68ºC 30s) with the exception of PDS 271-287, where the annealing temperature used was 58ºC. COBRA was performed with primers PDS 271-272 after bisulfite treatment via direction addition of 5U of *Bst*UI and digestion at 60ºC for 1 hour, followed by electrophoresis on a 4% agarose gel. In this assay, digestion products arise from methylated DNA; unmethylated DNA resists digestion.

### Targeted sequencing analysis

PCR products were pooled, column cleaned with the BioBasic EZ-10 DNA Columns (BioBasic BS427), eluted in 30 uL of Tris-HCl pH 8.0, and quantified via Nanodrop. NEBNext Universal Adaptors were ligated using T4 DNA Ligase in Quick Ligase Buffer in a total volume of 22 uL at 25□C for 15 minutes.

Self-ligated adaptors (adaptor dimer) was excluded with a 0.8X (2.55 uL) Agencourt AMPure XP bead cleanup (Beckman Coulter A63880) followed by 1X PEG-NACl buffer (25% PEG, 2.5 M NaCl in DEPC water) cleanup to exclude self-ligated adaptors. Barcodes were added via PCR amplification for 20 cycles with Phusion HF (NEB M0530L) using NEBNext Multiplex Oligos for Illumina® (NEB E7335). Adaptor dimer was again excluded via 0.8X AMPure bead cleanup followed by a 1X PEG Buffer-NaCl cleanup using the same beads. Libraries were quantified using Qubit, and sequenced on a MiSeq 2000 Paired End 2 × 250bp at the Cornell University Genomics Core. Samples were evaluated for quality and trimmed to 200bp with Trim Galore! (www.bioinformatics.babraham.ac.uk). Trimmed samples were probed for WT, *tm5.1*, or Igf2r DMR-specific sequences using the grep function; these reads were compiled into separate files and analyzed using QUMA [86]. Animals with a minimum of ten reads were included for analysis. Total reads per amplicon per sample are listed in Table S2. Grep sequences and reference sequences for QUMA are listed in Table S3.

## AUTHORS’ CONTRIBUTIONS

ETC, DHT, and PDS designed experiments. ETC, MH, and DHT performed experiments. ETC and PDS wrote the manuscript. All authors read and approved the final manuscript.

## ACKNOWLEDGEMENTS

We thank Roman Spektor for critical contributions to the design, collection, and reporting of these data. We thank Maria Garcia-Garcia for generous donation of *Sox2-Cre* mice.

We thank Bruce Morgan for providing the primer sequences for PDS 2794-5, which were used to genotype the *pA-TetOFF* and *TetOFF* alleles.

## ETHICAL APPROVAL AND CONSENT TO PARTICIPATE

All animal studies were approved by the Institutional Animal Care and Use Committee protocol 2002-0075. No studies included human subjects.

## SUPPORTING INFORMATION CAPTIONS

**Figure S1. Validation of *Rasgrf1*. a)** Generation of *Rasgrf1* from tm5.0PDS tm5.0PDS *Rasgrf1*. *Rasgrf1* (**a**, upper schematic) was generated via CRISPR-Cas9-mediated homology-directed repair in v6.5 embryonic stem cells. Targeting was confirmed by Sanger sequencing of PCR products generated with primers PDS 2359-8 and PDS 2344-2263, which respectively span the junctions of the 5′ and 3′ homologous arms of the pETC6 vector (grey boxes), and the flanking sequences of the *Rasgrf1*^tm5.0PDS^ allele. PDS 2344 falls within the neo resistance cassette (n). PDS 2358 sequence falls partially within TetO. DNAs from Wild-type (WT) animals show no product for either PDS 2359-8 or PDS 2344-2263. *Rasgrf1*^tm5.1 PDS^ (*tm5.1*, a, lower schematic) was generated by crossing *Rasgrf1*^tm5.0PDS^ males to females constitutively expressing FlpE recombinase (FlpE). Recombination was confirmed with Sanger sequencing of PCR products generated with primers PDS 2262-2263. PDS 2262 maps to TetO sequence, and the product contains the single residual frt site (orange triangle) remaining after recombination. PCR products shown below the amplicon schematics arise only when using DNAs from mutant animals. **b)** Schematic for Southern blot shown in **c)**; probe location, shown with pink line, is outside of the targeting vector; location of *Pst*I sites are indicated. Yellow triangles indicate the *Rasgrf1* repeats. Homologous arms are omitted for clarity. **c)** Southern blot of a *tm5.1* heterozygote vs. a wild-type animal *tm1* and a *Rasgrf1* homozygote (*tm1*); in these animals, the probe detects the same 3kb fragment in *tm1* as is *tm5.1*. **d)** Copy number qPCR support targeting of the endogenous allele: *tm5.1* heterozygotes have half the copy number of the repeat element as WT animals, but the same number of DMR sequences. Error bars represent standard error across biological duplicates.

**Figure S2. The *tm5.1* allele lacks imprinted methylation and expression in the absence of transactivator.** Summary of *Rasgrf1* expression, *Rasgrf1* imprinting, and total animals measured in *tm5.1* animals of the **a)** N2 and **b)** N3 generations. The majority of animals inheriting *tm5.1* paternally (+/*tm5.1*) expressed *Rasgrf1* biallelically as assayed by endpoint PCR, but levels were only 2% those seen in wild-type (WT) mice, and the alleles were unmethylated. However, a subset of +/*tm5.1* animals (4 of 10 in the N2, and 2 of 15 in the N3 generation) expressed *Rasgrf1* at WT levels from the paternal *tm5.1* allele. N3 animals expressing *Rasgrf1* paternally and at wild-type levels did not arise from methylated fathers, indicating that sporadic *tm5.1* DMR methylation, and *Rasgrf1* expression was not an inherited state. This is consistent with findings with the *Rasgrf1*^*tm1PDS*^ allele which lacks the *Rasgrf1* repeats [21]. **c)** Allele-specific expression analysis. A male chimera prepared using C57BL/6 blastocysts, and v6.5 ES cells with the *tm5.1* allele on the 129S4 (129) background, was crossed with C57BL/6 females. Neonatal brain cDNA was subjected to endpoint RT-PCR using primers spanning SNPs from the 129 and C57BL/6 backgrounds that harbor distinct *Aci*I sites. Product digestion with *Aci*I produces allele-specific bands, reporting the expressed allele(s). The slowest and fastest migrating bands represent the 129 paternal *Tm5.1* allele; the two middle bands represent the C57BL/6 allele. WT animals, inheriting the C57BL/6 paternal allele, expressed *Rasgrf1* solely from the WT C57BL/6 allele(s). A portion of +/*tm5.1* animals express paternally from the *tm5.1* 129 allele. The majority express biallelically from the maternal C57BL/6 and paternal 129 alleles. **d)** qRT-PCR of *Rasgrf1* in wild-type, and +/*tm5.1* animals. Paternally expressing +/5.1 animals express *Rasgrf1* at WT levels, whereas biallelically expressing +/5.1 animals express at 2% of WT, indicating biallelic expression detected in *tm5.1* by endpoint PCR was seen when the normally active paternal allele was silent. Error bars represent standard error across at least biological triplicate. **e)** Targeted bisulfite sequencing (bar graphs) and COBRA analyses (gel images) of the WT DMR in tail gDNA of WT animals (left), and the *tm5.1* DMR of +/*tm5.1* animals with paternal (middle) and biallelic (right) *Rasgrf1* expression. Animals with paternal expression were methylated at the *tm5.1* DMR, whereas animals with biallelic expression were hypomethylated. WT animals have two copies of the WT *Rasgrf1* DMR, one hypermethylated and one hypomethylated. As such, bisulfite analysis of the WT DMR in soma is 50%. Bar graphs report the percentage of total reads with the levels of methylation shown in the key on the right. COBRA queries methylation at five *Bst*UI sites in both the WT and the *tm5.1* DMRs. “+” and “–” denote addition or lack of *Bst*UI. Methylated (+mC) and unmethylated –mC) sites are respectively sensitive or resistant to *Bst*UI digestion. Digestion products are indicated by black arrowheads. The PCR product of the WT DMR shows partial digestion, reporting the different methylation states of the two parental alleles; paternally expressing +/*tm5.1* show full digestion of the *tm5.1* DMR; biallelically expressing +/*tm5.1* animals show no digestion. **f)** +/*tm5.1* animals are lower in body weight compared to their WT littermates at several ages post-weaning, consistent with previous findings that loss of paternal methylation and expression at *Rasgrf1* leads to diminished body weight [41]; four to ten mice per genotype at each age were measured.

**Figure S3. Generation of the *TetOFF* allele.** *pA-TetOFF* males were crossed with females carrying a *Sox2-Cre* transgene to generate the recombined *TetOFF* allele. PDS 2794 maps to the splice acceptor site of the ROSA26 locus; PDS 2795 maps 3’ of the *TetOFF* coding sequence, within wild-type ROSA26 sequence. pA-*TetOFF* animals produce a 2.2kb PCR product with PDS 2794-5, whereas *TetOFF* animals produce a 1.5kb PCR product due to loss of the floxed Neo-polyA cassette. Note that a portion of animals are mosaic for pA-*TetOFF* and *TetOFF* (mosaic); though only fully recombined mice bearing the *TetOFF* allele were analyzed in crosses with the *tm5.1* allele.

**Figure S4. Successful pitRNA induction in *TetON*/*tm5.1* and *TetOFF*/*tm5.1* several tissues. a)** Endpoint RT-PCR for pitRNA in the tissues of transactivated *tm5.1* 28 to 42 day old animals demonstrate induction of pitRNA in several tissues of *TetON: +*/*tm5.1* and *TetOFF: +*/*tm5.1* mice. Low signals from testes are likely due to the blood testes barrier restriction entry of doxycycline (dox) in adult males. pitRNA is not detectable by endpoint PCR in wild-type (WT) animals. **b)** qPCR for pitRNA in adult liver shows 10-fold upregulation of pitRNA in *TetON*/*tm5.1* animals fed dox chow, and 500-fold upregulation of pitRNA in *TetOFF*/*tm5.1* livers relative to WT. **c)** qPCR for pitRNA in neonatal brain shows nearly 10,000 fold upregulation of pitRNA in *TetOFF*/*tm5.1* brains relative to *WT*, *TetOFF/+*, and +/*tm5.1* animals. ***, p < 10^−6^; n.s., not significant.

**Figure S5. Additional bisulfite analysis of embryonic and adult male germline. a)** COBRA of the *tm5.1* DMR for the *TetOFF/tm5.1* animal depicted in **Fig 2a** (#1) as well as a littermate of the same genotype (#2). “+” and “–” designate addition of *Bst*UI. The *tm5.1* DMR is hypomethylated in the gonocyte and somatic fractions of both animals. Targeted bisulfite sequencing for #2 is shown below. The paternally inherited wild-type (WT) DMR is hypermethylated in both gonocyte and somatic fractions; the *tm5.1* DMR is hypomethylated in both gonocyte and somatic fractions. The Igf2r DMR is hypomethylated in gonocyte and 50% methylated in soma. **b)** COBRA for the *tm5.1* DMR for the *TetON/tm5.1* + dox animal depicted in **Fig 2c** (#1) as well as a littermate of the same genotype (#2). Bisulfite sequencing results for #2 are depicted below. Note that while the *tm5.1* DMR is hypomethylated in the gonocyte and somatic fractions of #1, it is approximately 50% methylated in #2 by bisulfite sequencing and COBRA. This is consistent with rates of stochastic *tm5.1* DMR methylation as previously described: As described in **Fig S2**, 10-25% of *+/tm5.1* show evidence of *tm5.1* methylation and expression in the soma. Rep #2 is one of six assayed (18%) transactivated *tm5.1* embryonic gonads to display partial methylation of the *tm5.1* DMR in the gonocyte and somatic fraction. **c)** COBRA for the two *+/tm5.1* and two *TetOFF/tm5.1* animals shown in **Fig 2e** (#1 and #2 of each genotype). COBRA results for an additional three *+/tm5.1* animals (#3, #4, #5) are shown; bisulfite results for #3 are below. COBRA from an additional *TetOFF/tm5.1* animal (#3) is also shown. In all samples, the *tm5.1* DMR is hypomethylated. **Bisulfite** Key at lower right.

**Figure S6. Transactivation of *tm5.1* with *TetON* and doxycycline induces expression of *Rasgrf1* from the *tm5.1* allele, but does not impart methylation to the *tm5.1* DMR or affect expression or methylation of the WT allele. a)** *Aci*I digestion of PDS 245-6 endpoint RT-PCR product in the offspring of a *TetON* x *tm5.1* cross. *TetON*: +/*tm5.1* animal express from the paternal *tm5.1* (129) allele, whereas WT and *TetON*:+/+ animals express from the paternal WT (C57BL/6) allele, and +/*tm5.1* animals express weakly and biallelically. **b)** *Aci*I digestion of PDS 245-6 endpoint RT-PCR product in the offspring of a *tm5.1* x *TetON* cross. *TetON*: *tm5.1*/+ animals express *Rasgrf1* biallelically, indicating activation of the maternal *tm5.1* allele in addition to the normally active paternal WT allele. *tm5.1*/+, *TetON*: +/+ and WT animals express paternally from the WT allele. **c)** COBRA for the *tm5.1* and WT DMRs in tail DNA of *TetOFF*: +/*tm5.1* and *TetOFF*: *tm5.1*/+ animals. The *tm5.1* DMR is hypomethylated regardless of parental origin. The WT DMR is unmethylated if inherited maternally (animal at left) but fully methylated, as evidenced by complete digestion of PCR products, if inherited paternally (animal at right).

**Figure S7. Oocyte preloading of pitRNA to 90X wildtype levels has no effect on *Rasgrf1* expression in wild-type offspring. a)** pitRNA levels in oocytes after induction in *TetON:* +/*tm5.1* females treated with doxycycline. Error bars represent standard error across biological duplicates **b)** RT-PCR for *Zp3*, an oocyte-specific transcript, was used to confirm isolation of oocytes. **c)** The pedigree shown was used to determine if oocyte loading of pitRNA could produce intergenerational activation of the maternal *Rasgrf1* allele. pitRNA was induced in oocytes of a *TetON: +*/*tm5.1* female by IP injection of doxycycline for three days prior to mating; doxycycline-containing chow was provided throughout pregnancy. Activation of the maternal allele in neonatal brains of offspring was used to report intergenerational effects, with allele-specific expression assayed by *Aci*I digestion of RT-PCR products. All mice tested expressed only the paternal allele. **d)** *Rasgrf1* levels in brains of animals depicted directly above in c). Error bars represent standard error across biological duplicates at minimum. **e)** COBRA of the wild-type allele in WT, *TetON*: +/+, and *tm5.1*/+ tails from three representative animals from d) (purple trapezoids). The WT DMR is partially methylated in WT and *TetON*: +/+ animals (having inherited two WT DMRs), and fully methylated in the *tm5.1*/+ animal (having inherited the WT DMR paternally); the *tm5.1* DMR is hypomethylated in this animal. Pale horizontal bar in the top third of each panel represents the dye front. ** p < 0.01; n.s., not significant; IP, intraperitoneal.

**Table S1. Primer sequences for all analyses described.** All qPCR primers are between 87 and 113% efficient. Unless otherwise noted, all endpoint PCRs were performed using Promega GoTaq in 25 uL reactions, Ta = 60C, te = 30s for 35 cycles. All qPCRs are performed in 20 uL reactions using SYBR Green Master Mix (ABI 4367659) on a Biosystems 7500 with annealing temperature 60ºC for forty cycles followed by a dissociation stage.

**Table S2. Total reads per DMR by Sample ID.** Amplicons for which less than ten reads were recovered were excluded from analysis. Total number of reads correlates in part to total amplicons for each barcode (variable). n.a., not applicable; n.m., not measured.

**Table S3. Sequences used for QUMA.** In preparation for analysis, raw files were probed for reads containing amplicon-specific sequences using the grep function in Linux. QUMA was performed on these reads.

**Table S4. Total reads broken down by C57 and FVB fractions for MiSeq sequencing of PDS 245-6 RT-PCR product in neonatal brain**. The *tm5.1* allele was generated on a 129 background; *Rasgrf1* expressed from the *tm5.1* allele carries 129 SNPs. *TetON* and *TetOFF* are carried on a C57Bl/6 (B6) background; “+” always designates a B6 allele. A B6 x FVB gDNA sample is sequenced as a control. n.i., not included in paper.

**Table S5. SNP IDs and grep sequences for allele-specific PDS 245-6 digestion and sequencing**. SNPs were probed for using the grep function and quantified using the wc function in Linux.

